# NRT1.1-dependent NH_4_^+^ toxicity in *Arabidopsis* is associated with disturbed balance between NH_4_^+^ uptake and assimilation

**DOI:** 10.1101/241174

**Authors:** Shaofen Jian, Qiong Liao, Haixing Song, Qiang Liu, Joe Eugene Lepo, Chunyun Guan, Jianhua Zhang, Zhenhua Zhang

## Abstract

A high concentration of a sole ammonium (NH_4_^+^) source in growth media is often toxic to plants. The nitrate transporter NRT1.1 is involved in plant NH_4_^+^ toxicity; however, its mechanism remains undefined. In this study, wild-type *Arabidopsis* (Col-0) and *NRT1.1* mutants (*chl1-1* and *chl1-5*) were grown hydroponically in NH_4_NO_3_ and (NH_4_)_2_SO_4_ media to evaluate NRT1.1 function in NH_4_^+^ stress responses. All plants grew normally in mixed N sources, but Col-0 displayed more chlorosis, and lower biomass and photosynthesis than the *NRT1.1* mutants in the (NH_4_)_2_SO_4_ condition. Grafting experiments between Col-0 and *chl1-5* further confirmed that NH_4_^+^ toxicity is NRT1.1-dependent. In (NH_4_)_2_SO_4_ medium, NRT1.1 facilitated the higher expression of NH_4_^+^ transporters, increasing NH_4_^+^ uptake. Additionally, glutamine synthetase (GS) and glutamate synthetase (GOGAT) in roots of Col-0 plants decreased and soluble sugar accumulated significantly, whereas pyruvate kinase (PK)-mediated glycolysis was not affected, all of which contributed to NH_4_^+^ accumulation. In contrast, the *NRT1.1* mutants reduced NH_4_^+^ accumulation and enhanced NH_4_^+^ assimilation through glutamate dehydrogenase (GDH) and glutamate-oxaloacetate transamination (GOT) activity. In addition, the upregulation of genes involved in senescence in Col-0 plants treated with (NH_4_)_2_SO_4_ suggests that ethylene could be involved in NH_4_^+^ toxicity responses. Our results indicate that NH_4_^+^ toxicity is dependent on NRT1.1 in *Arabidopsis*, characterized by enhanced NH_4_^+^ accumulation and by perturbed NH_4_^+^ metabolism, which stimulated ethylene-induced plant senescence.

**Highlight:** Nitrate transporter NRT1.1 enhances NH_4_^+^ accumulation, disturbs the NH_4_^+^ metabolism, and aggravates NH_4_^+^ toxicity in *Arabidopsis* when grown under sole NH_4_^+^ condition.

## Introduction

Ammonium (NH_4_^+^) is a paradoxical inorganic nitrogen source in the soil that is available to plants during growth and development. It is the preferable N source for some species such as rice (*Oryza sativa* L.) and tea (*Camellia sinensis*) (Gao et al., 2010, Ruan et al., 2016). However, the high levels of NH_4_^+^ in the soil due to increased N fertilizer application, for example, is detrimental to higher terrestrial plants and can promote chlorosis, necrosis, and decreased biomass and yield (Chen et al., 2013; Li et al., 2013). To date, the primary underlying mechanisms of these forms of NH_4_^+^ toxicity in plants remain unclear (Li et al., 2013).

Excess NH_4_^+^ uptake and accumulation have been proposed to contribute to NH_4_^+^ toxicity in plants. Acidification of the rhizosphere from proton excretion during NH_4_^+^ uptake is considered to be the principal contributor to NH_4_^+^ toxicity (Chaillou et al., 1991); however, some studies suggest that the initial step of NH_4_^+^ toxicity is pH-independent (Britto and Kronzucker, 2002; van den Berg et al., 2005).

NH_4_^+^ serves as the ubiquitous intermediate metabolite in N metabolism in plants (Joy, 1988; Glass et al., 1997). Its high concentration in plant cells is damaging to plant tissues. NH_4_^+^ accumulation is commonly observed under high NH_4_^+^ conditions. In rice (*Oryza sativa* L.), the application of high levels of NH_4_^+^ resulted in NH_4_^+^ accumulation and futile NH_4_^+^ cycling (Chen et al., 2013). NH_4_^+^ accumulation occurs in part from NH_4_^+^ uptake. In *Arabidopsis thaliana*, the ammonium transporter (AMT) family consists of six protein isoforms, three of which are AtAMT1;1, AtAMT1;2, and AtAMT1;3 and are located in the root where they are responsible for about 90% of NH_4_^+^ uptake (Yuan et al., 2007a). Wang et al. (2013) verified that under high-NH_4_^+^ stress (30 mM NH_4_^+^), AtAMT1;3 is endocytosed from the plasma membrane to reduce NH_4_^+^ uptake and mitigate NH_4_^+^ toxicity. In addition, the capacity to detoxify excessive NH_4_^+^ accumulation in plant cells is thought to be an important metabolic factor for alleviating the consequence of this stress (Hoai et al., 2003). Glutamine synthetase (GS) acts in the center of NH_4_^+^ flow in ammonium metabolism (Miflin and Habash, 2002), and glutamate dehydrogenase (GDH) is an anti-stress enzyme involved in ammonia detoxification (Skopelitis et al., 2006). It has been reported that NH_4_^+^-tolerant plants have greater GS activity and less NH_4_^+^ accumulation in plant tissues (Cruz et al., 2006; Omari et al., 2010). Konishi et al. (2017) reported that *GLN1* plays a dominant role in regulating NH_4_^+^ uptake in *Arabidopsis*. Indeed, pharmacological studies suggest that Gln rather than NH_4_^+^ could be the real signaling molecule that regulates the expression of NH_4_^+^ transport and assimilation genes (Tabuchi et al., 2007). It is suggested that NH_4_^+^ uptake and assimilation are integrated; nevertheless, the coordination between NH_4_^+^ uptake and assimilation during NH_4_^+^ toxicity remains unclear.

Ethylene serves as an intermediate for signal transduction and translation, and manipulates the physiological responses under stress (Ismail and Horie, 2017; Arraes et al., 2015; Zhang et al., 2014; Li et al., 2014; Cao et al., 2007). Conversely, severe stress causes disequilibrium in ethylene production and accelerates plant senescence (Djanaguiraman and Vara, 2010; Jing et al., 2005). The occurrence of stress symptoms frequently accompanies ethylene evolution and NH_4_^+^ accumulation under drought, salt stress, and NH_4_^+^ stress (Barker and Corey, 1991; Feng and Barker, 1992). By exogenous application of NH_4_^+^ to leaves, Li et al. (2013) observed a significant increase in ethylene production and the inhibition of lateral root initiation in *Arabidopsis*. However, the relationship between NH_4_^+^ accumulation and ethylene evolution/production, as well as their contribution to stress induced symptoms differs between plant species. Lin et al. (2002) suggest that NH_4_^+^ accumulation in detached rice leaves does not change tissue sensitivity to ethylene, and ethylene is not involved in NH_4_^+^-promoted senescence; however, the specifics of how ethylene could be involved in the development of NH_4_^+^ toxicity symptoms remain to be elucidated.

*NRT1.1* is the primary member of the nitrate transporter (*NRT*) gene family in higher plants (Tsay et al., 1993). It has multiple functions in physiological processes in plants such as NO_3_^−^ and auxin transport, NO_3_^−^ signal sensing, and stomatal movement (Tsay et al., 1993; Guo et al., 2003; Krouk G. et al., 2010; Bouguyon et al., 2015). Many studies have revealed that NRT1.1 is essential for plants to defend themselves against unfavorable environments, including cadmium stress, salt stress, and proton toxicity (Mao et al., 2014; Aragón and Navarro, 2017; Fang et al., 2016). Interestingly, the *Arabidopsis NRT1.1* mutants *chl1-1* and *chl1-5* display higher resistance to high concentrations (10 mM) of sole NH_4_^+^ sources than that of its wild-type plants (Hachiy et al., 2011; Hachiy and Noguch, 2011). However, the mechanism for how NRT1.1 mediates NH_4_^+^ uptake and the subsequent emergence of NH_4_^+^ toxicity still need to be identified at physiological and molecular levels.

Here, we analyzed the gene expression, metabolites, and the physiological and chemical activities in roots and shoots of wild type and *NRT1.1* mutant *Arabidopsis* plants grown in nutrient growth media that contained high concentrations of NH_4_^+^. We aimed to clarify: 1) whether NH_4_^+^ toxicity symptoms are NRT1.1-dependent; 2) the physiological role of NH_4_^+^ uptake and assimilation during NH_4_^+^ toxicity; and 3) the function of ethylene in NRT1.1-mediated NH_4_^+^ toxicity. Our results provide further insight into the understanding of NH_4_^+^ toxicity in plants.

## Materials and methods

### Experimental materials and growth conditions

Seeds of wild-type *Arabidopsis* (Col-0), *NRT1.1* knockout mutants (*chl1-1*, *chl1-5*), an *NRT1.1* point mutation (*chl1-9*) line, and an ethylene signal mutant (*ein2*) were sown in nutritional soil media in a growth chamber (300 μmol photons·m^−2^·s^−1^, 16 h photoperiod, 22 °C) and allowed to germinate and grow for 10 d; then, seedlings with one pair of true leaves were transplanted to 600-mL pots and cultivated hydroponically with nutritional media. The growth media contained 1.125 mM NH_4_NO_3_, 1.25 mM KCl, 0.625 mM KH_2_PO_4_, 0.5 mM MgSO_4_, 0.5 mM CaCl_2_, 1.25 μM Fe-EDTA, 17.5 μM H_3_BO_3_, 3.5 μM MnCl_2_, 0.25 μM ZnSO_4_, 0.05 μM NaMoO_4_, and 0.125 μM CuSO_4_. The pH of the nutrient media was adjusted to 6.0 and MES (2.5 mM) was added to the nutrient media to buffer any changes in the pH. The media was refreshed every four days. There were 9 plants in each pot and the growth condition was the same as mentioned above. After 4 weeks, half of the plants were transferred to 5 mM (NH_4_)_2_SO_4_ medium for an additional 10 d. The control group was supplied with 5 mM NH_4_NO_3_. The experiment was arranged in a completely randomized design. The position of the pots was interchanged when refreshing the solution to eliminate any edge effects.

### ^15^N pulse assay

To investigate the uptake of NH_4_^+^ during (NH_4_)_2_SO_4_ treatment, Col-0, *chl1-1*, and *chl1-5* seedlings were hydroponically cultivated for 4 weeks and then divided into two groups for 5% atom abundance labeling (^15^NH_4_)_2_SO_4_ (pH 6.0) at the start of the sole NH_4_^+^ treatment and 5 d after the treatment, respectively. Before ^15^N labeling, roots were washed with 0.1 mM CaSO_4_ for 1 min and then placed in 5 mM (^15^NH_4_)_2_SO_4_ medium for 5 days. When sampling, the roots were washed again with 0.1 mM CaSO_4_ for 1 min and then with deionized water. Root and shoot tissues were harvested separately and oven-dried at 70 °C for 48 h. The samples were ground into powder with TissueLyser-48 (Jingxin Co. Ltd., China) and ^15^N abundance in samples was determined using a continuous-flow isotope ratio mass spectrometer coupled with a carbon-nitrogen elemental analyzer (ANCA-MS; PDZ Europa).

### Grafting experiment

A grafting experiment was performed as described in Rus et al. (2006) using an SZX-ILLB200 microscope (OLYMPUS OPTICAL CO., LTD, Japan). Col-0 and *chl1-5* seeds were sterilized in 75% ethanol and germinated on ½ MS plates containing 3 mg·L^−1^ benomyl, 0.04 mg·L^−1^ benzyladenine, 0.02 mg·L^−1^ indole-3-acetic acid, and 17 g·L^−1^ agar. All plates with stratified seeds were placed vertically in a growth chamber as described above. Five-day-old seedlings were grafted on the plate with a MANI Ophthalmic Knife (Straight 15, SWISSMED, Japan). The grafted seedlings were placed on plates for another 5 d until the graft union formed. Grafted seedlings without adventitious roots were cultivated hydroponically and treated with (NH_4_)_2_SO_4_ as described.

### Ethylene regulation experiment

Four-week-old Col-0, *chl1-1*, and *chl1-5* seedlings were divided into three groups for (NH_4_)_2_SO_4_ treatment. The leaves of the first and second groups were treated with 5 μM aminoethoxyvinylglycine (AVG) and 50 μM 1-aminocyclopropanecarboxylic acid (ACC), respectively, at 08:00 every day throughout the (NH_4_)_2_SO_4_ treatment course. The third group was sprayed with deionized water and served as a control. Each plant was sprayed until the leaves were thoroughly wetted. After 10 d of treatment, chlorophyll content was determined and pictures were taken to determine the phenotype.

### Photosynthesis measurements

Photosynthesis of the youngest fully expanded leaves in rosettes was measured 10 d after (NH_4_)_2_SO_4_ treatment using an LI-6400 portable photosynthesis system (Li-Cor Inc., Lincoln, NE, USA). Air temperature in the cuvette during measurement was maintained at 22 °C and the photosynthetic photon flux intensity (PPFD) was 200 μmol·m^−2^·s^−1^. The CO_2_ concentration in the cuvette was adjusted to 500 μmol·mol^−1^ with a CO_2_ mixer and the VPD was at 1.0–1.5 kPa. Data were recorded after equilibration to a steady state. After photosynthetic determination, the leaf area was measured with a leaf area meter (CI-202 Portable Laser Area Meter, CID Bio Science Inc, USA). The data for gas exchange parameters were adjusted to the accurate leaf areas.

### Measurement of chlorophyll concentration

Chlorophyll in fresh rosette leaves of NH_4_NO_3_ (control) and (NH_4_)_2_SO_4_-treated plants was extracted according to the method reported by Wellburn and Lichtenthaler (1984). The absorbance of the extract was measured at 663 nm and 645 nm using a UV-VIS spectrophotometer (UV-2600, Shimadzu, Japan) to estimate total chlorophyll concentrations.

### Measurements of biomass, total nitrogen content, and NH_4_^+^ and soluble sugar concentrations

Plants in (NH_4_)_2_SO_4_ treatment and control groups were harvested and the roots and shoots were separated. The samples were oven-dried at 70 °C to a constant weight. Dry samples were ground into a fine powder and about 100 mg of each sample were digested with H_2_SO_4_-H_2_O_2_ at 350 °C for N quantification. Total N was determined with an AA3 Autoanalyzer (SEAL, Germany).

NH_4_^+^ and soluble sugar concentrations in fresh samples were extracted with deionized water. NH_4_^+^ concentrations were measured using the method of indophenol blue colorimetry at 630 nm (Zanini, 2001) and (NH_4_)_2_SO_4_ was used as a standard. Soluble sugar concentrations were determined by the anthrone-sulfuric acid method (Wang et al., 2002).

### Amino acid quantification

Amino acid components in root and shoot of NH_4_NO_3_ and (NH_4_)_2_SO_4_-treated plants were quantified with HPLC. The samples were frozen immediately in liquid N_2_ and stored at -80 °C. Frozen leaf samples (200 mg) were powdered with liquid N_2_, and homogenized with 1.5 mL 0.1% phenol in 6 M HCl. The homogenates were hydrolyzed for 22 h at 100 °C. After cooling, 1 mL hydrolysate was dried with organomation (NDK200-2, Hangzhou Mi’ou Instrument Co. Ltd. China) and re-dissolved with 1 mL 0.1 M HCl. For derivation of amino acids, 200 μL re-dissolved hydrolysate was mixed with 20 μL norleucine internal standard solution, 200 μL triethylamine acetonitrile (pH > 7), and 100 μL isothiocyanate acetonitrile and placed at 25 °C for 1 h. Then 400 μL hexane was added and samples were incubated for 10 min with shaking. The underlayer solution was filtered with 0.45 μm syringe filter. The analyses were performed on the Rigol L3000 (Rigol L3000, Beijing RIGOL Technology Co., Ltd. China). Chromatographic separation was operated in RP-HPLC ACE column (5C18-HL), particle size 5 μm (250 mm × 4.6 mm), at circumstances temperature through the binary gradient. Mobile phase A contain 25 mM acetate buffer (pH 6.5) and 70 mL acetonitrile. Mobile phase B was 80% acetonitrile aqueous solution. The flow rate was 1.0 mL·min^−1^ and column temperature was 40 °C. The sample was maintained for 45 min.

### Enzyme activity assays

Frozen samples (100 mg) were ground into powder in liquid N_2_, and homogenized with 3 mL 50 mM Tris-HCl buffer (pH 8.0) containing 2 mM Mg^2+^, 2 mM DTT, and 0.4 M sucrose. The homogenate was centrifuged at 12×*g* for 10 min at 4 °C. The supernatant was used to determine the activities of the N metabolic enzymes glutamine synthase (GS), NADH-dependent glutamate synthase (NADH-GOGAT), NADH-dependent glutamate dehydrogenase (NADH-GDH), glutamic-oxaloacetic transaminase (GOT), and glutamic-pyruvic transaminase (GPT).

GS and GOGAT activities were measured according to the method described previously by Zhang et al. (1997) and Singh and Srivastava (1986), respectively. Activity of NADH-GDH was determined according to the method described by Loulakakis and Roubelakis-Angelakis (1990). GOT and GPT activities were assayed using the method described by Wu et al. (1998).

Frozen samples were pulverized in liquid N_2_, and about 100 mg of each sample were used to measure pyruvate kinase (PK) activity according to Lepper et al. (2010) with some modification. The samples were extracted with 3 mL 100 mM Tris-HCl (pH 7.5) containing 10 mM β-mercaptoethanol, 12.5% (v/v) glycerine, l mM EDTA-Na_2_, 10 mM MgCl_2_, and 1% (m/v) PVP-40. These solutions were centrifuged at 12×*g* and 4 °C for 10 min. The reaction mixture contained 100 mM Tris-HCl (pH 7.5), 10 mM MgCl_2_, 0.16 mM NADH, 75 mM KCl, 5.0 mM ADP, 7.0 units L-lactate dehydrogenase (LDH), and 1.0 mM phosphoenolpyruvate (PEP). The reaction was started by adding 0.1 mL enzyme extract to 0.9 mL of the reaction mixture, and was carried out at 25 °C to monitor the changes of absorbance at 340 nm for 180 s.

Enzyme activity was expressed as moles of metabolite generated/consumed per mg protein per unit of time. Soluble protein content was measured using the Bradford method with BSA as the standard (Bradford, 1976).

### RNA extraction and RT-PCR assay

Seedlings were treated with 5 mM NH_4_NO_3_ and (NH_4_)_2_SO_4_ for 3 d before the roots and shoots were harvested separately for RNA analysis. Total RNA was extracted with TRIzol (Invitrogen, USA), precipitated with an equal volume of isopropanol, washed with 75% ethanol, and dissolved with RNase-free water. cDNAs were synthesized using the PrimeScript™ RT Kit with gDNA Eraser (Perfect Real Time) (TAKARA, Japan) following the manufacturer’s protocol. The relative expression of genes in roots and shoots was determined by quantitative RT-PCR performed on an Applied Biosystems StepOne™ Real-Time PCR System using SYBR Premix Ex-Taq (TAKARA) according to the manufacturer’s protocol. Primes used in the assays are listed in **Supplementary Table 1**, and the expression data were normalized to *Actin2*.

### Statistics

The experiment was conducted according to a completely random design. Four biological replicates and two technical replicates were provided for each treatment. Two-way analysis of variance (ANOVA) was conducted to analyze the effects of N source and plant material. Multiple comparisons were performed using the least significant difference (*LSD*) multiple range test. The differences between control and (NH_4_)_2_SO_4_ treatments in the same plant material were evaluated with the Student’s *t*-test. Differences were considered statistically significant at a *P* < 0.05 level.

## Results

### NH_4_^+^ toxicity is NRT1.1-dependent

Compared to control, a high concentration of a sole NH_4_^+^ source supplied in the form of (NH_4_)_2_ succinate, (NH_4_)_2_SO_4_, or NH_4_Cl, respectively, for 10 days caused chlorosis in the leaves in Col-0, with the most serious chlorosis observed in plants treated with (NH_4_)_2_SO_4_ (**Fig. 1A**). However, the leaf color of *chl1-1* and *chl1-5* mutant plants was relatively less affected by sole NH_4_^+^ source treatment. In this study, we used (NH_4_)_2_SO_4_ treatment for all subsequent experimentation and analysis.

**Fig. 1.**
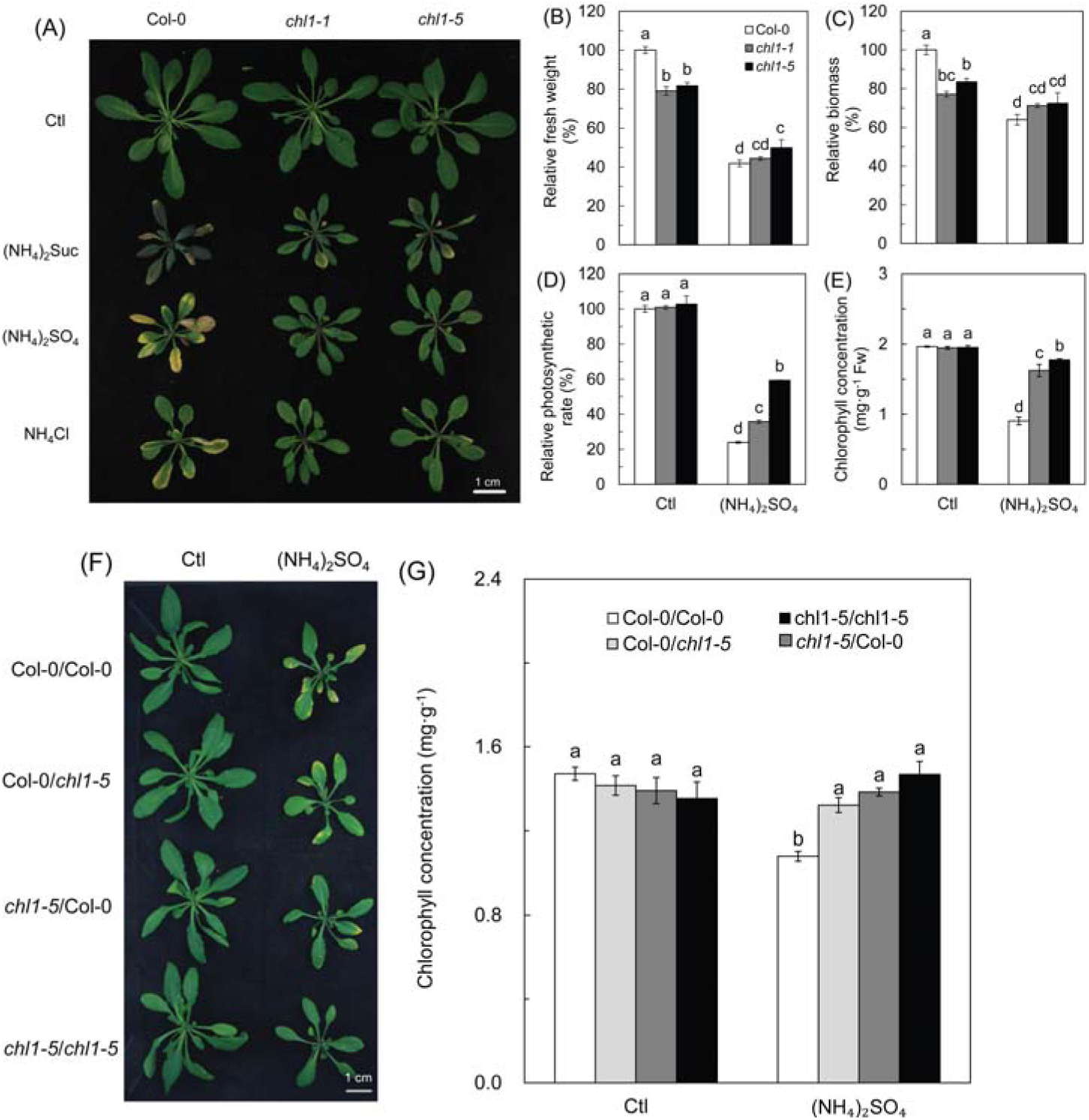
Effects of high concentrations of sole NH_4_^+^ sources on *Arabidopsis* growth and photosynthesis **(A)** Plants were grown in 5 mM NH_4_NO_3_ (control) and different sources of ammonium (10 mM) media for 10 d. **(B)** Relative fresh weight, **(C)** biomass, **(D)** photosynthetic rates, and **(E)** chlorophyll content of Col-0, *chl1-1*, and *chl1-5* grown in 5 mM NH_4_NO_3_ (control) or (NH_4_)_2_SO_4_ for 10 d. **(F)** Grafted plants grown in 5 mM NH_4_NO_3_ (control) and (NH_4_)_2_SO_4_ for 10 d. Plant material names on the left of indicate the genotypes of the shoot/root combination. **(G)** Chlorophyll content of the grafted plants grown in 5 mM NH_4_NO_3_ (control) and (NH_4_)_2_SO_4_ for 10 d. The relative values were calculated with the average value of Col-0 in control conditions as a reference. Fresh weight and dry weight were obtained from pooled samples of seven plants from each replicate. Data represent means ± SE (*n* = 4). Bars with the same letter indicate no significant difference at *α* = 0.05 using the *LSD* method. Scale bars in (A) and (F) represent 1 cm.

Application of a sole NH_4_^+^ source reduced the plant size of all plant materials dramatically (**Fig. 1A**). The fresh weight of *chl1-1* and *chl1-5* was reduced by 44% and 39%, and biomass was lowered by 10% and 16%, respectively, compared to the 58% and 36% reduction in fresh weight and biomass, respectively, observed for Col-0 (**Fig. 1B and C**). In contrast to *chl1-1* and *chl1-5*, the phenotype of *chl1-9* was consistent with Col-0 (**Fig. S1A and B**) when grown in (NH_4_)_2_SO_4_ medium. There was no difference in photosynthetic rates and chlorophyll between Col-0 and *NRT1.1* mutants in NH_4_NO_3_ growth conditions, whereas (NH_4_)_2_SO_4_ reduced chlorophyll and photosynthetic rates in all genotypes with a larger reduction measured in Col-0 than in *chl1-1* and *chl1-5* (**Fig. 1D and E**). Nevertheless, there was no significant difference in stomatal conductance between these genotypes under (NH_4_)_2_SO_4_ treatment (**Fig. S1C**). These results indicate that NRT1.1 could be involved in NH_4_^+^ toxicity in *Arabidopsis*.

To further confirm that NH_4_^+^ toxicity is NRT1.1-dependent, we established grafted Col-0 and *chl1-5* plants that could be grown under NH_4_^+^ conditions. As shown in **Fig. 1F and G**, the plants with Col-0 roots and/or shoots developed chlorosis under (NH_4_)_2_SO_4_ conditions. Plants with Col-0 shoots displayed worse chlorosis than those with Col-0 roots. Taken together, the results indicate that NH_4_^+^ toxicity is NRT1.1-dependent, but independent of the functions of *NRT1.1* in signaling and stomatal regulation (**Fig. S1**).

### NRT1.1 increases NH_4_^+^ uptake and accumulation in roots and shoots

Total N content, NH_4_^+^ concentration, and the expression of *AMT1* genes in response to NH_4_^+^ transport were assayed to identify the difference in NH_4_^+^ uptake between Col-0 and *NRT1.1* mutants. As shown in **Fig. 2A**, there was no difference among Col-0, *chl1-1*, and *chl1-5* plants in total N before (NH_4_)_2_SO_4_ treatment. In contrast, treatment with (NH_4_)_2_SO_4_ for 10 d decreased total N significantly in the three genotypes, with the exception of Col-0 roots; the total N in Col-0 was significantly higher than that measured in *chl1-1* and *chl1-5* **(Fig. 2B)**. There was no significant difference in NH_4_^+^ concentration in the three genotypes under control conditions **(Fig. 2C)**. (NH_4_)_2_SO_4_ treatment significantly reduced NH_4_^+^ concentrations in *chl1-1* and *chl1-5* roots and shoots, while the NH_4_^+^ concentration in Col-0 was stable in the root and significantly increased in the shoot (**Fig. 2C**). Similarly, the relative ^15^N atom abundance dramatically decreased with (NH_4_)_2_SO_4_ in the roots of *chl1-1* and *chl1-5*, but increased in shoots of Col-0 plants (**Fig. 2D**). Although the relative ^15^N atom abundance in *chl1-1* and *chl1-5* shoots significantly increased under (NH_4_)_2_SO_4_ treatment, the increase was much smaller than that measured in Col-0 (**Fig. 2D**).

**Fig. 2.**
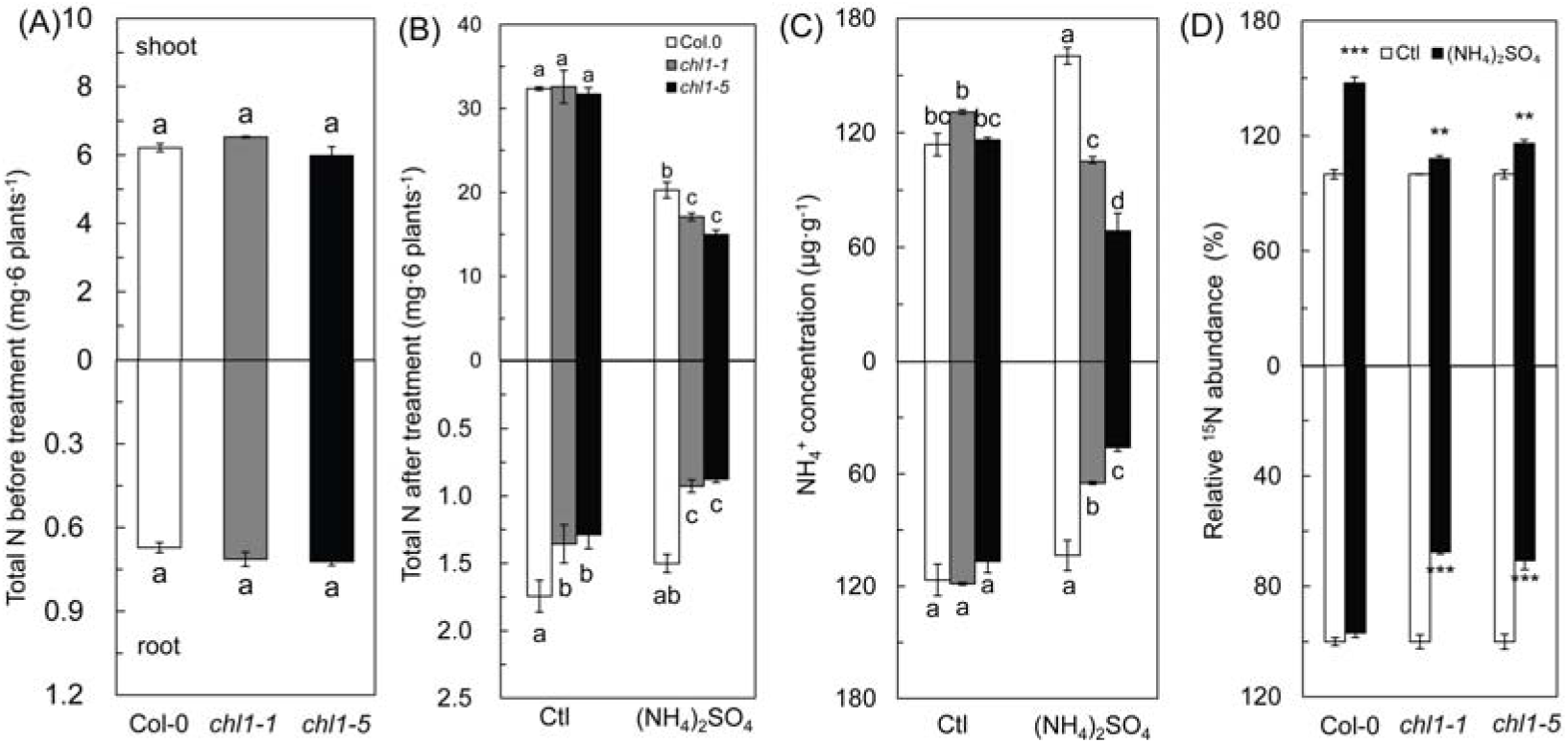
Effects of a high sole NH_4_^+^ source on N accumulation and NH_4_^+^ uptake in *Arabidopsis*. **(A)** Total N content of Col-0, *chl1-1*, and *chl1-5* plants grown in 1.125 mM NH_4_NO_3_ medium before (NH_4_)_2_SO_4_ treatment. **(B)** Total N content and **(C)** NH_4_^+^ concentration of Col-0, *chl1-1*, and *chl1-5* plants grown in 5 mM NH_4_NO_3_ (control) and (NH_4_)_2_SO_4_ for 10 d. **(D)** Relative ^15^N abundance (%) in Col-0, *chl1-1*, and *chl1-5* plants grown in 5 mM NH_4_NO_3_ (control) and (NH_4_)_2_SO_4_ for 5 d. Total nitrogen content was obtained from pooled samples of seven plants from each replicate. Data represent means ± SE (*n* = 4). Bars with the same letter are not significantly different at *α* = 0.05 by the *LSD* method. Bars with two (**) and three (***) asterisks indicate significant differences from the control at *α* = 0.01 and *α* = 0.001, respectively, using a two-tailed Student’s *t*-test.

The expression of *AMT1;1*, *AMT1;2*, *AMT1;3*, and *AMT1;5* in *chl1-1* and *chl1-5* was not significantly different from that in Col-0 under control condition, whereas *AMT1;3*, and *AMT1;5* in Col-0 were up-regulated when treated with (NH_4_)_2_SO_4_, especially *AMT1;5*, which was expressed 200-fold higher than that in the control **(Fig. 3C-D)**. The expression level of *AMT1* genes in *chl1-1* and *chl1-5* was significantly lower (*AMT1;1*) or unchanged (*AMT1;2* and *AMT1;3*) under (NH_4_)_2_SO_4_ with respect to that of control **(Fig. 3A-D)**. The expression of *AMT1;5* was significantly higher in all genotypes in response to (NH_4_)_2_SO_4_; however, the expression in Col-0 was higher than that in *NRT1.1* mutants (**Fig. 3D**). With (NH_4_)_2_SO_4_ treatment, the expression of all *AMT1* genes in Col-0 was significantly higher than that in *chl1-1* and *chl1-5* **(Fig. 3A-D)**. These results suggest that *NRT1.1* facilitates the expression of *AMT1*s in *Arabidopsis* in response to (NH_4_)_2_SO_4_, and ultimately results in increased NH_4_^+^ uptake.

**Fig. 3.**
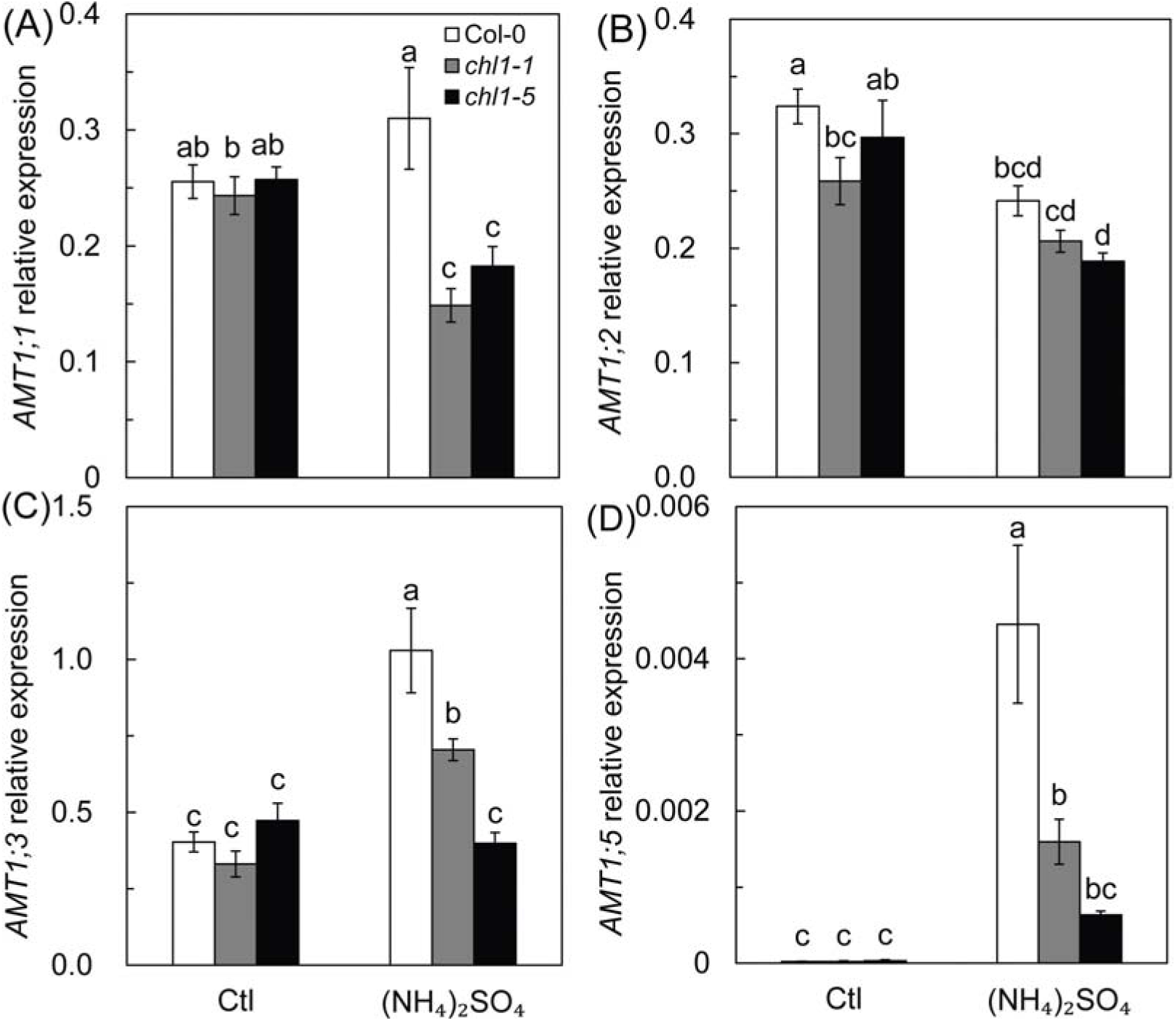
*AMT1* gene expression under high sole NH_4_^+^ source treatment in roots of Col-0, *chl1-1*, and *chl1-5* plants grown in 5 mM NH_4_NO_3_ (control) and (NH_4_)_2_SO_4_ for 3 d. Roots were sampled for the expression of **(A)** *AMT1;1* **(B)** *AMT1;2* **(C)** *AMT1;3* and **(D)** *AMT1;5*. Data represent means ± SE (*n* = 4). Bars with the same letter are not significantly different at *α* = 0.05 using the *LSD* method.

### *NRT1.1* inhibits the activity of NH_4_^+^ metabolism enzymes under high concentration of sole NH_4_^+^ condition

Under control conditions, GS activity was significantly lower in *chl1-1* and *chl1-5* than in Col-0 in roots **(Fig. 4A)**. The GS activity was decreased in Col-0 plants treated with (NH_4_)_2_SO_4_, but was not affected in *chl1-1* and *chl1-5* under the same growth conditions **(Fig. 4A)**. In shoots, (NH_4_)_2_SO_4_ significantly increased GS activity in Col-0 and *chl1-1* but not in *chl1-5* **(Fig. S3A)**.

**Fig. 4.**
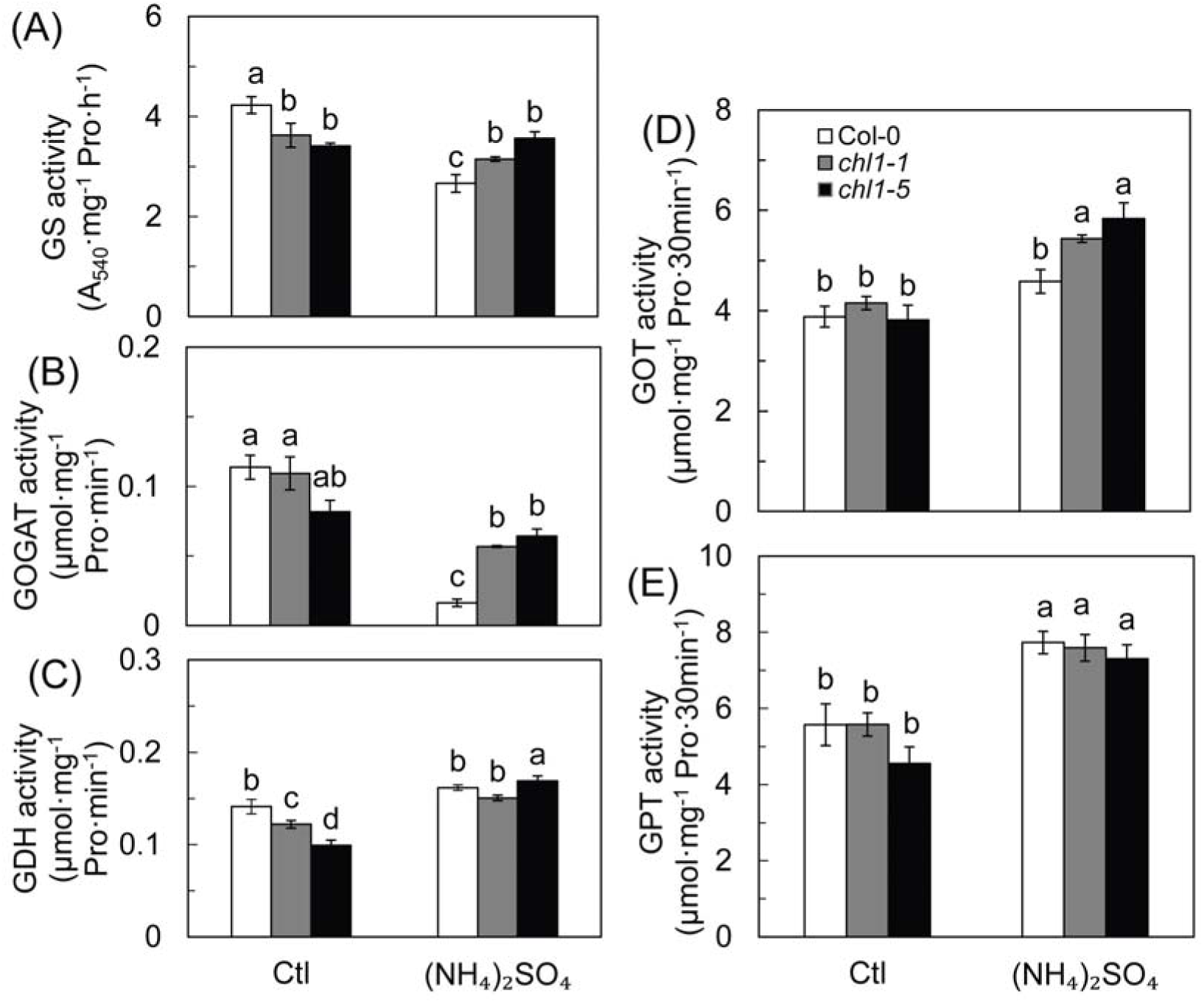
Activity of NH_4_^+^ assimilation enzymes and transaminases under high sole NH_4_^+^ source treatment in roots of Col-0, *chl1-1*, and *chl1-5* plants grown for 10 d in 5 mM NH_4_NO_3_ (control) and (NH_4_)_2_SO_4_. **(A)** Glutamine synthetase (GS), **(B)** glutamate synthetase (GOGAT), **(C)** glutamate dehydrogenase (GDH), **(D)** glutamate-oxaloacetate transaminase (GOT), and **(E)** glutamate-pyruvic transaminase (GPT) activity in roots of Col-0, *chl1-1*, *chl1-5* plants. Data represent means ± SE (*n* = 4). Bars with the same letter are not significantly different at *α* = 0.05 by the *LSD* method.

GOGAT activity in roots did not differ between genotypes in control **(Fig. 4B)**. (NH_4_)_2_SO_4_ significantly reduced GOGAT activity by 85.64% in Col-0, whereas activity in *chl1-1* and *chl1-5* decreased by 48.13%, and 21.54%, respectively **(Fig. 4B)**. In shoots, (NH_4_)_2_SO_4_ significantly increased GOGAT activity in *chl1-1* and *chl1-5* but not in Col-0, whereas no difference was observed in control conditions among the genotypes **(Fig. S3B)**.

GDH activity was significantly higher in Col-0 than in *chl1-1* and *chl1-5* in root under control conditions **(Fig. 4C)**. (NH_4_)_2_SO_4_ had no significant influence on GDH activity in Col-0, but increased GDH activity in *chl1-1* and *chl1-5* **(Fig. 4C)**. In shoots, (NH_4_)_2_SO_4_ increased GDH activity in all genotypes, with no differences among genotypes under control or (NH_4_)_2_SO_4_ treatments **(Fig. S3C)**.

There were no differences in GOT and GPT activities in roots of the three genotypes under control conditions **(Fig. 4D-E)**; however, (NH_4_)_2_SO_4_ increased GOT activity in *chl1-1* and *chl1-5* but not in Col-0 **(Fig. 4D)**. Although GPT activity in the three genotypes was higher under (NH_4_)_2_SO_4_ treatment than in the control, there was no significant difference between the genotypes **(Fig. 4E)**. In shoots, GOT and GPT activities were not affected by (NH_4_)_2_SO_4_ treatment, and no difference in their activities could be measured among the wild type and mutant genotypes **(Fig. S3D-E)**. Taken together, these results indicate that *NRT1.1* has a negative effect on the activities of NH_4_^+^ assimilation enzymes and transaminases in response to a high concentration of a sole NH_4_^+^ source.

### NRT1.1 disturbs the balance between carbon and NH_4_^+^ metabolism in sole NH_4_^+^ source conditions

Soluble protein concentrations in *chl1-1* and *chl1-5* were significantly higher than those in Col-0 under control conditions with the exception of *chl1-1* roots, which showed lower soluble protein concentrations than the wild type in the control treatment (**Fig. 5A**). Irrespective of shoot or root, (NH_4_)_2_SO_4_ increased soluble protein concentrations in Col-0, but not in *chl1-1* and *chl1-5*; while the concentrations in shoots did not differ between the genotypes; mutants showed lower concentrations in roots than did Col-0 (**Fig. 5A**).

**Fig. 5.**
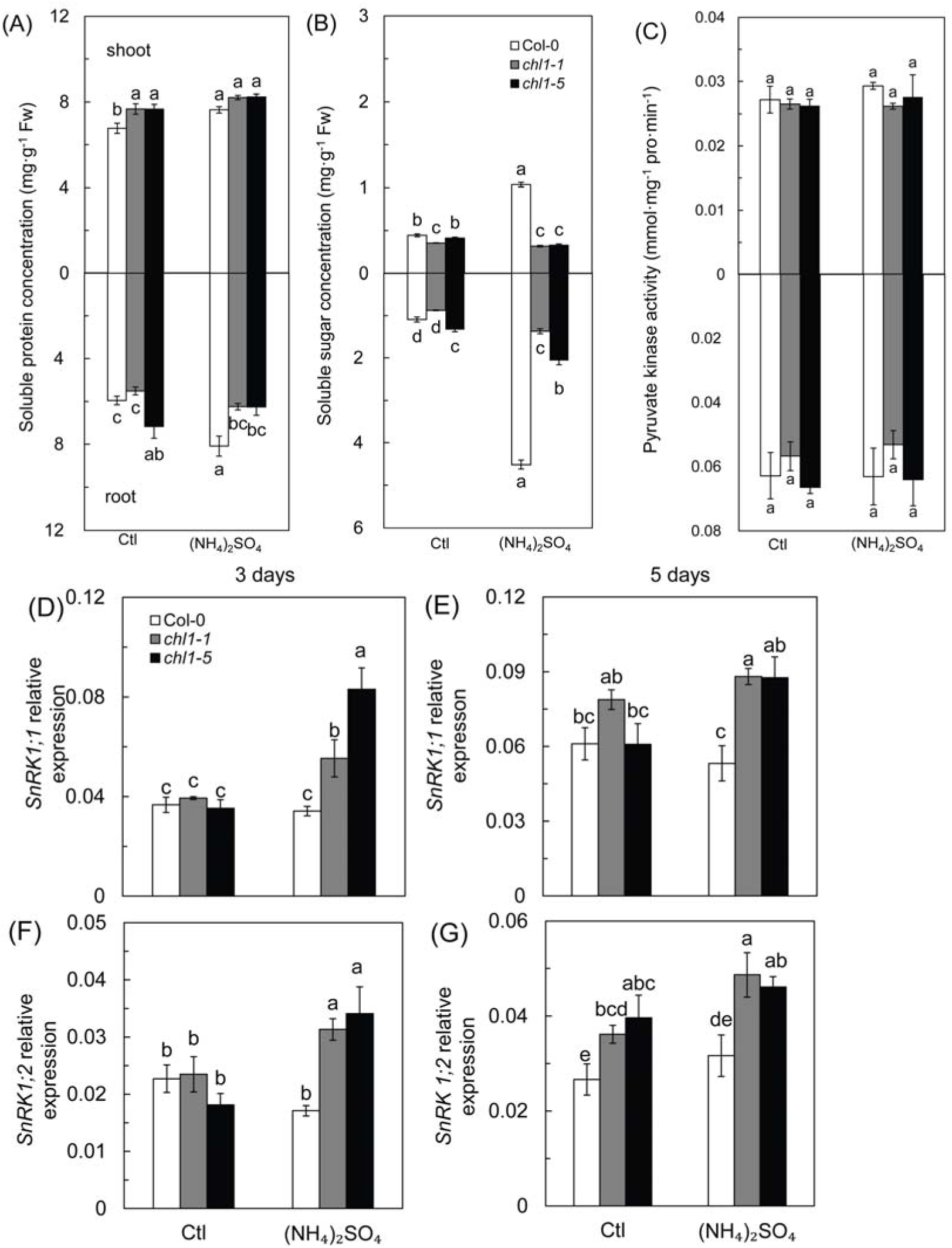
Soluble protein concentration **(A)**, soluble sugar content **(B)**, activity of pyruvate kinase **(C)** and expression of *SnRK1* genes with high sole NH_4_^+^ source treatment in Col-0, *chl1-1*, and *chl1-5* plants **(D-G)**. *SnRK1;1* and *SnRK1;2* expression after 3 d **(D** and **F)** and 5 d **(E** and **G)** of (NH_4_)_2_SO_4_ treatment is shown. Plants measured in **A-C** were grown in 5 mM NH_4_NO_3_ (control) and (NH_4_)_2_SO_4_ media for 10 d before the roots and shoots were harvested. Plants measured in **D-G** was grown in 5 mM NH_4_NO_3_ (control) and (NH_4_)_2_SO_4_ medium for 3 and 5 d, respectively, before mRNA was extracted from roots. Data represent means ± SE (*n* = 4). Bars with the same letter are not significantly different at *α* = 0.05 by the *LSD* method.

Soluble sugar concentrations in roots and shoots varied slightly in the wild type and mutants in the control treatment **(Fig. 5B)**; whereas the soluble sugar concentrations were 4.14 and 2.33 times higher in roots and shoots of Col-0, respectively, under (NH_4_)_2_SO_4_ conditions than under control conditions **(Fig. 5B)**. However, soluble sugar concentrations in *chl1-1* and *chl1-5* were not as affected by (NH_4_)_2_SO_4_, and were significantly lower than that in Col-0 **(Fig. 5B)**. Pyruvate kinase (PK) activity was consistent among genotypes regardless of (NH_4_)_2_SO_4_ treatment (**Fig. 5C**).

The gene expression of the carbon and nitrogen metabolic regulator *SnRK1* was studied to evaluate the coordination of carbon and nitrogen metabolism. As shown in **Fig. 5D-G**, after 3 d in control conditions, the expression of *SnRK1;1* and *SnRK1;2* was unchanged in wild type and mutant lines; however, *SnRK1;2* expression increased after 5 d in the mutants compared to wild type. With (NH_4_)_2_SO_4_ treatment for 3 d, we observed an up-regulation of these two genes in the mutants, but their expression in Col-0 was unchanged (**Fig. 5D and F**). After 5 d of treatment, *NRT1.1* mutants expressed higher levels of the *SnRK1* genes than Col-0 with (NH_4_)_2_SO_4_ treatment (**Fig. 5E and G**).

To further investigate the effect of (NH_4_)_2_SO_4_ on nitrogen metabolism, we compared the difference in amino acid composition between Col-0 and *chl1-1* grown with different N sources. As shown in **Fig. 6A-D**, major amino acids such as aspartate (Asp) and asparagine (Asn) were less affected by (NH_4_)_2_SO_4_ in roots than in shoots in wild type and mutant lines **(Fig. 6A and B)**. Glutamate (Glu) and glutamine (Gln) in roots of *chl1-1* were significantly lower than that in Col-0 with (NH_4_)_2_SO_4_ treatment **(Fig. 6C and D)**. In shoots, there were no significant differences in amino acid content between Col-0 and *chl1-1* under control conditions **(Fig. 6A-D)**; however, (NH_4_)_2_SO_4_ treatment increased the amino acid content with the exception of Asn in Col-0, which was lower than that measured in control conditions **(Fig. 6A-D)**. In shoots, Asp, Asn, and Gln, but not Glu, were significantly higher in *chl1-1* than in Col-0 under (NH_4_)_2_SO_4_ treatment **(Fig. 6A-D)**. The ratio of amino acids in root and shoot were calculated and **Fig. 6E-H** indicates that in control conditions, there was no noticeable change in amino acid ratios between the wild type and mutants. (NH_4_)_2_SO_4_ promoted a higher accumulation of Asn in Col-0 roots than in shoots, whereas the R:S ratio of Asn in *chl1-1* was not significantly affected **(Fig. 6F)**. The R:S ratio of Gln dramatically declined with (NH_4_)_2_SO_4_ treatment in both Col-0 and *chl1-1*; however, no difference was observed between these lines (**Fig. 6H**).

**Fig. 6.**
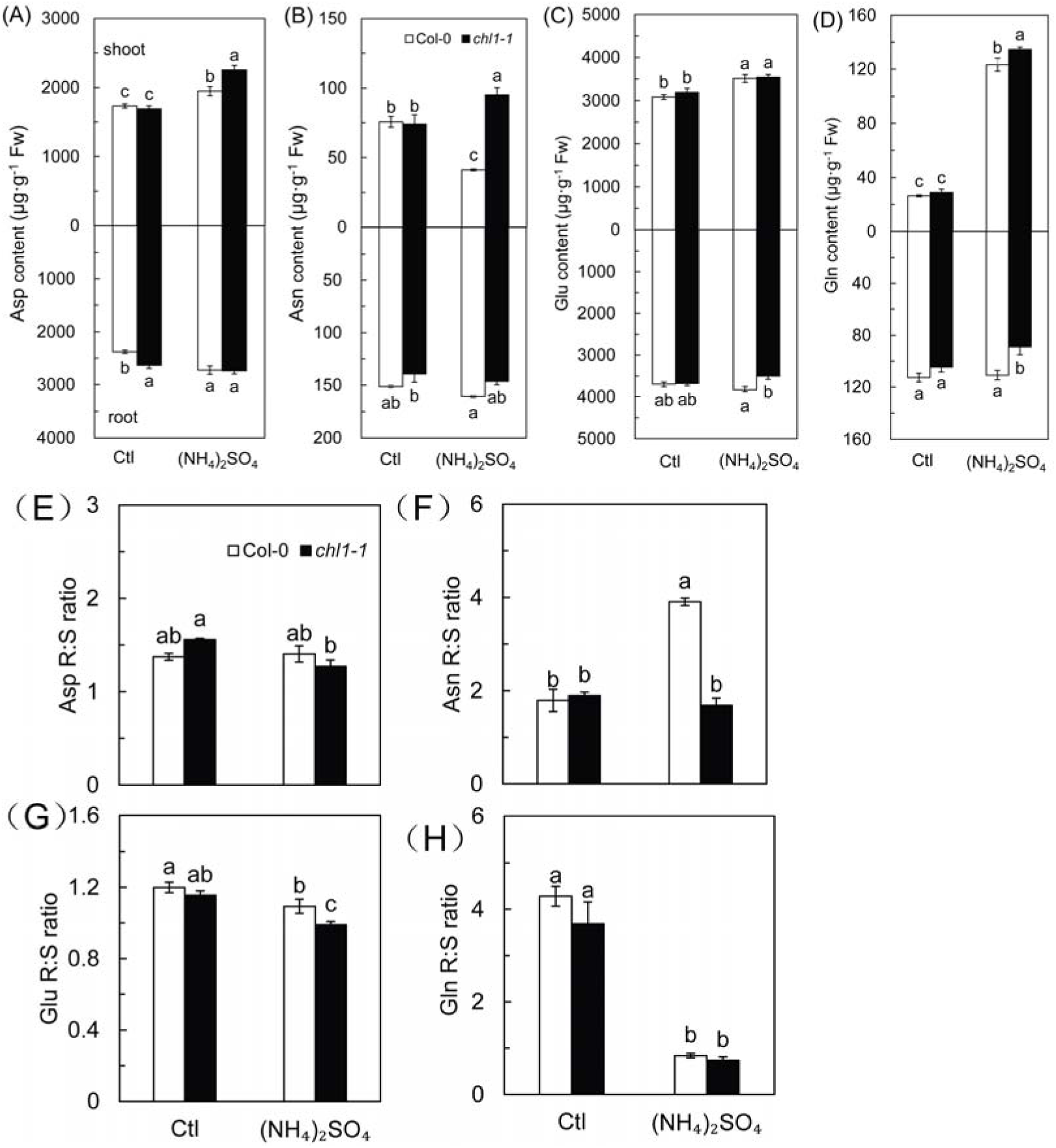
Free-amino acid content and their R:S ratios with high concentrations of a sole NH_4_^+^ source in Col-0 and *chl1-1*. Plants were grown in 5 mM NH_4_NO_3_ (control) and (NH_4_)_2_SO_4_ media for 10 d. Asp **(A)**, Asn **(B)**, Glu **(C)**, and Gln **(D)** was measured in the shoots and roots, and then the root/shoot ratios of Asp **(E)**, Asn **(F)**, Glu **(G)**, and Gln **(H)** were calculated. Data represent means ± SE (*n* = 3). Bars with the same letter are not significantly different at *α* = 0.05 according to the *LSD* method.

### *NRT1.1*-dependent NH_4_^+^ toxicity is associated with ethylene-induced plant senescence

Using ethylene mutant *ein2*, and foliar application of the ethylene precursor 1-aminocyclopropanecarboxylic acid (ACC) and ethylene antagonist aminoethoxyvinylglycine (AVG) in wild type and *NRT1.1* mutants, we tested whether NRT1.1-dependent NH_4_^+^ toxicity is associated with ethylene evolution. As shown in **Fig. 7A**, the growth of *ein2* plants under (NH_4_)_2_SO_4_ treatment did not differ from that of the Col-0; furthermore, it maintained a higher chlorophyll concentration under these conditions as well (**Fig. 7B**). While (NH_4_)_2_SO_4_ treatment significantly increased NH_4_^+^ concentration in *ein2* and Col-0, there was no significant difference between the two lines (**Fig. 7C**).

**Fig. 7.**
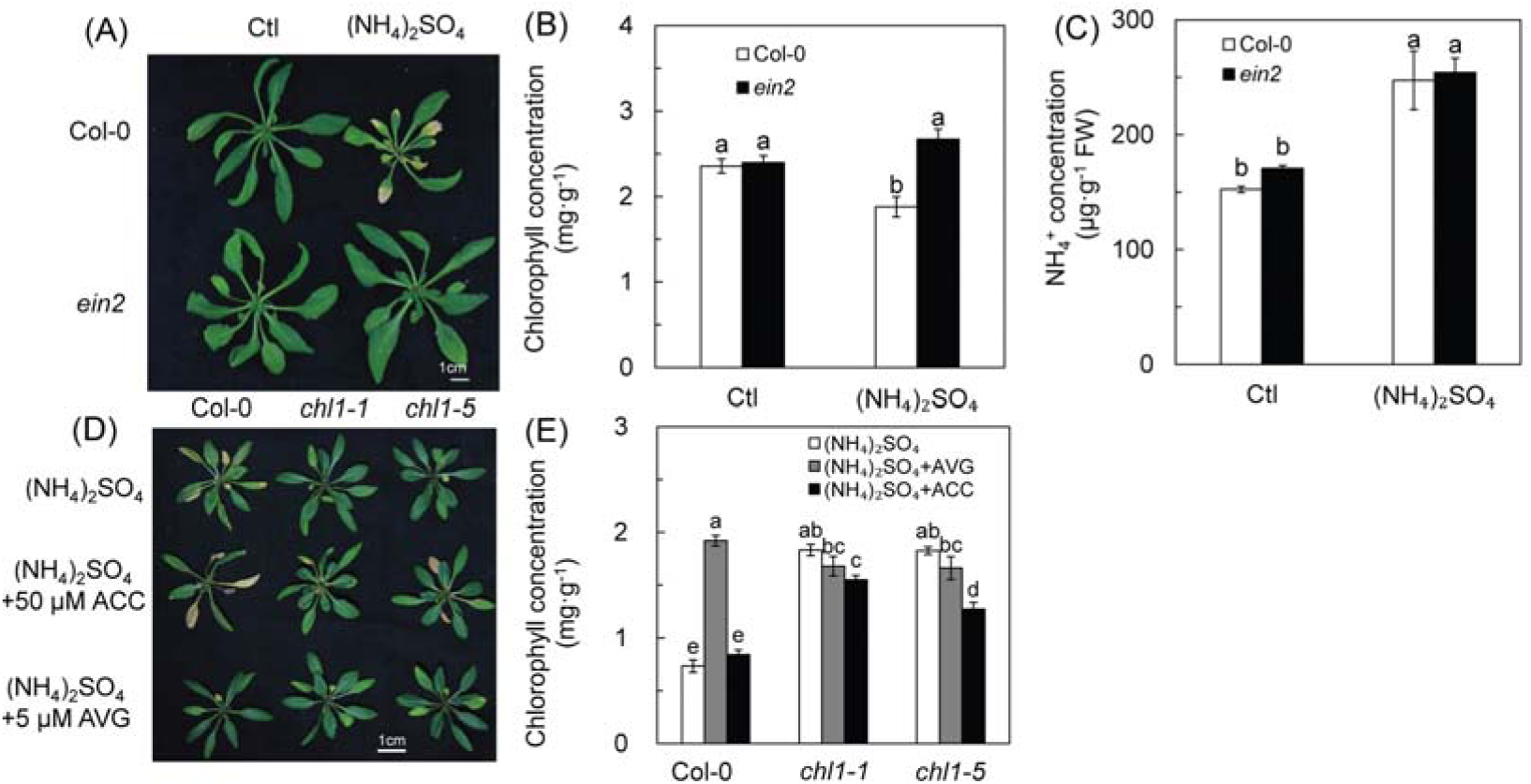
Ethylene is involved in NRT1.1-dependent NH_4_^+^ toxicity **(A)** Col-0 and the ethylene mutant *ein2* were grown in 5 mM NH_4_NO_3_ (control) and (NH_4_)_2_SO_4_ media for 10 d. **(B)** Chlorophyll content and **(C)** NH_4_^+^ concentration of Col-0 and *ein2* plants grown in 5 mM NH_4_NO_3_ (control) and (NH_4_)_2_SO_4_ media for 10 d. **(D)** Col-0, *chl1-1*, and *chl1-5* were grown in 5 mM (NH_4_)_2_SO_4_ media and then their leaves were sprayed with 50 μM ACC or 5 μM AVG, respectively, at 08:00 for 10 d throughout the treatment period. The control was sprayed with deionized water. **(E)** Chlorophyll content of Col-0, *chl1-1* and *chl1-5* plants grown in 5 mM (NH_4_)_2_SO_4_ medium and treated with ACC and AVG, respectively, for 10 d. Data represent means ± SE (*n* = 4). Bars with the same letter are not significantly different at *α* = 0.05 by the *LSD* method.

Applying AVG alleviated the chlorosis in Col-0 after (NH_4_)_2_SO_4_ treatment, whereas *chl1-1* and *chl1-5* were unchanged (**Fig. 7D and E**). ACC application caused similar levels of chlorosis with (NH_4_)_2_SO_4_ treatment in Col-0, and the chlorophyll content in *chl1-1* and *chl1-5* significantly decreased with ACC treatment in the presence of (NH_4_)_2_SO_4_ (**Fig. 7D and E**).

The expression of the senescence-related genes *WRKY45*, *WRKY53*, *SAG13*, *SAG29*, *SAG113*, and *GEN4* were assayed after treating Col-0, *chl1-1*, and *chl1-5* with (NH_4_)_2_SO_4_ **(Fig. S5A-F)**. All genes were significantly up-regulated by (NH_4_)_2_SO_4_ treatment in Col-0; in contrast, the expression of these genes in *chl1-1* and *chl1-5* was less up-regulated or even down-regulated, as was the case for *GEN4* **(Fig. S5A-F)**. These results suggest that NRT1.1-dependent NH_4_^+^ toxicity is associated with ethylene and its induction of plant senescence.

## Discussion

### NRT1.1-dependent symptoms of NH_4_^+^ toxicity

Chlorosis and a decline in biomass are the most typical phenotypes in plants grown in high concentration of sole NH_4_^+^ sources (Chen et al., 2013). These phenotypes are consistent with the reduced photosynthetic rates and chlorophyll concentrations that were observed in *Arabidopsis* wild-type plants in this study **(Fig. 1 A-E)**. Similar to the results reported by Hachiya and Noguchi (2011), the *NRT1.1* knockout mutants *chl1-1* and *chl1-5* had a stronger resistance to NH_4_^+^ toxicity relative to Col-0, had weaker chlorosis, and less reduction in biomass, photosynthetic rate, and chlorophyll content (**Fig. 1 A-E**). These results indicated that *NRT1.1* is responsible for the NH_4_^+^ toxicity in *Arabidopsis* when grown under high concentrations of a sole NH_4_^+^ source. Our grafting experiment further demonstrated that NH_4_^+^ toxicity in *Arabidopsis* is *NRT1.1*-dependent regardless of its expressing in roots or shoots (**Fig. 1 F and G**).

*NRT1.1* is a multifunctional gene that participates in plant development and the interaction between plants and their environment (Bouguyon et al., 2015). In addition to being an NO_3_^−^ transporter, NRT1.1 serves as NO_3_^−^ sensor in roots and is also a stomatal regulator in guard cells (Guo et al., 2003), which are tightly correlated with plant growth and physiological responses to environment. The *chl1-9* mutant is defective in nitrate uptake but shows a normal primary nitrate response (Ho et al., 2009) and showed NH_4_^+^ toxicity symptoms similar to that in Col-0 in response to (NH_4_)_2_SO_4_ (**Fig. S1A and B**), suggesting a specific deficiency in NO_3_^−^transport capacity is not enough to cause NH_4_^+^ toxicity in *Arabidopsis* in the absence of NO_3_^−^. As the phosphorylation state and amount of NRT1.1 protein are similar in both wild-type and *chl1-9* (Hachiya and Noguchi, 2011), we speculate that the altered phosphorylation state of NRT1.1 and the interaction between NRT1.1 and other proteins in metabolic pathways could relate to the occurrence of NH_4_^+^ toxicity. The performance of *chl1-9* in response to (NH_4_)_2_SO_4_ in this study was different from that reported by Hachiya and Noguchi (2011); this discrepancy could be due to the variation in sensitivity to NH_4_^+^ at different growth stages. *NRT1.1* expression in guard cells mediates the accumulation of NO_3_^−^ during stomatal opening (Guo et al., 2003). Stomatal closure increases the resistance of CO_2_ diffusion from the ambient air into the intercellular space of leaves, and consequently limits the photosynthesis (Lawlor and Tezara, 2009). In this study, stomatal conductance (*g*_s_) in Col-0, *chl1-1*, and *chl1-5* displayed no significant difference with (NH_4_)_2_SO_4_ treatment (**Fig. S1C**). These results reveal that NRT1.1-dependent NH_4_^+^ toxicity in *Arabidopsis* is independent of the function of NRT1.1 in NO_3_^−^ transport and stomatal regulation.

### NRT1.1-related perturbation of carbon and nitrogen metabolism under high concentration of a sole NH_4_^+^ source

The interaction between carbon (C) and nitrogen (N) metabolism is of paramount importance to improve stress tolerance of plants (Reguera et al., 2013). The assimilation of NH_4_^+^ requires carbohydrates to provide the carbon skeletons of 2-oxoglutarate (2-OG) generated from the tricarboxylic acid (TCA) cycle (Yuan et al., 2007b). Nguyen et al. (2005) proposed that a reduction in photosynthetic capacity and decrease in 2-OG production might be responsible for the excess accumulation of NH_4_^+^. The availability of sugar in plant tissues may regulate N (including NH_4_^+^) uptake and assimilation, but depends on a series of carbon metabolic steps downstream of hexose phosphorylation (Lejay et al., 2003). Pyruvate kinase (PK) is at a crucial position to control the glycolysis of assimilated C for providing energy and carbon skeletons (2-OG) for N metabolism (Stitt, 1999). In this study, the activity of PK was not significantly different between Col-0 and *NRT1.1* mutants or between the control and experimental treatments, suggesting the availability of C skeletons is not a limiting factor in N assimilation, and that C and N metabolism in Col-0 were uncoupled when the plants were subjected to high concentrations of a sole NH_4_^+^ source. Complex regulatory machineries are involved in the coordination between N and C metabolism, including sugar, nitrate, and amino acid sensing, as well as phytohormones (Nunes-Nesi, et al., 2010). SnRK1 (SNF1-related protein kinase) plays a vital role in plant energy homeostasis, growth and development, and stress tolerance (Baenagonzález et al., 2007), where it regulates the key enzymes in C and N metabolism. The up-regulated expression of *SnRK1;1* and *SnRK1;2* in *NRT1.1* mutants were consistent with the increased N metabolism and resistance to NH_4_^+^ toxicity, suggesting *SnRK1* is an essential factor in C and N metabolism under NH_4_^+^ stress **(Fig. 5D-G)**.

Accumulation of NH_4_^+^ was often companied by the accumulation of free amino acids (Hoai et al., 2003), probably due to the lower requirement of NH_4_^+^ for amino acid synthesis. In this study, however, *chl1-1* had a higher amino acid content but lower NH_4_^+^ in (NH_4_)_2_SO_4_ conditions. This could have resulted from a greater NH_4_^+^ assimilation capacity and amino acid metabolism **(Fig. 4, Fig. 6, and Fig. S4**). Glutamine (Gln) and asparagine (Asn) are important for organic N storage and transport in plant tissues. They function as important signals that can reflect the N status in plants and mediate NH_4_^+^ uptake and assimilation (Tabuchi et al., 2007; Konishi et al., 2017). There was less Gln and Asn accumulation in roots in *chl1-1* than in Col-0 during (NH_4_)_2_SO_4_ treatment **(Fig. 6E and F)**. This potentially benefited the *chl1-1* mutant because less root Gln and Asn accumulation might increase the requirement for NH_4_^+^ incorporation into *de novo* amino acids, and help to mitigate NH_4_^+^ accumulation.

Increased amino acid synthesis could be advantageous to physiological processes such as stress defense and the accumulation of precursors for biosynthetic enzymes (Joshi et al., 2010). Glu can be converted into Asp and Ala by glutamic-oxaloacetic transaminase (GOT) and glutamic-pyruvic transaminase (GPT), respectively (Brock et al., 1970), and subsequently used for the synthesis of branched-chain amino acids (BCAAs). BCAAs such as Met, Thr, and Ile are derived from Asp pathway and provide precursors for many plant secondary metabolites, which could improve the resistance of plants to stresses (Joshi et al., 2010). In this study, *chl1-1* and *chl1-5* showed enhanced GOT activity under (NH_4_)_2_SO_4_ conditions (**Fig. 4D**). Furthermore, this was consistent with the increase in Ile and Thr contents (**Fig. S4**). Our results indicate that the Asp-derived pathway for amino acid synthesis is of great significance for improving the NH_4_^+^ toxicity tolerance in plants.

### NRT1.1 induces NH_4_^+^ accumulation in tissues under high sole NH_4_^+^ source medium and facilitates ethylene synthesis and plant senescence

Excessive NH_4_^+^ accumulation is the major cause for injury of plants in many unfavorable conditions (Barker, 1999; Hoai et al., 2003; Nguyen et al., 2005). In this study, NH_4_^+^ significantly accumulated in Col-0 under (NH_4_)_2_SO_4_ conditions **(Fig. 2C)**. The NH_4_^+^ equilibrium in plant cells is determined by NH_4_^+^ uptake, transport, and assimilation (Nguyen et al., 2005). The results obtained from our measurement of an ^15^N tracer revealed that NH_4_^+^ uptake was substantially higher in Col-0 than in mutants grown in sole NH_4_^+^ source conditions, and the absorbed NH_4_^+^ was mainly distributed in shoots **(Fig. 2D)**. This result indicates that increased NH_4_^+^ uptake in Col-0 made a significant contribution to NH_4_^+^ accumulation under such NH_4_^+^ conditions. *AtAMT1;1*, *AtAMT1;2*, and *AtAMT1;3* are the dominant genes required for NH_4_^+^ uptake in *Arabidopsis* (Yuan et al., 2007a). The up-expression of *AMT1* genes further suggests that they are regulated by NRT1.1 and account for the higher NH_4_^+^ accumulation in Col-0 than that in the *NRT1.1* mutants **(Fig. 3 A, B, and D)**.

Proton secretion is accompanied with NH_4_^+^ uptake and leads to acidification of the growth medium. Acidification of media has severely negative effects on *Arabidopsis* growth (Fang et al., 2016). It has been suggested that NH_4_^+^ uptake induces reduction in pH in the rhizosphere, which is one of the major effects of NH_4_^+^ toxicity (Chaillou et al., 1991). Present data excluded the effect of medium acidification on NH_4_^+^ toxicity, as the symptoms of NH_4_^+^ toxicity appeared only in Col-0, whereas the pH was not significantly different among the genotypes evaluated (**Fig. S2**).

Because stress generally causes NH_4_^+^ accumulation in plant cells, a high capacity to ameliorate NH_4_^+^ toxicity has been viewed as an important factor in alleviating the consequence of this stress (Hoai et al., 2003; Nguyen et al., 2005). The GS-GOGAT cycle continuously provides the substrates glutamate (Glu) for incorporation of NH_4_^+^ into glutamine (Gln) to detoxify excess NH_4_^+^ (Bittsánszky et al., 2015). The reduced activities of GS and GOGAT in Col-0 under the (NH_4_)_2_SO_4_ conditions in this study indicated that attenuated NH_4_^+^ assimilation capacity contributed to the accumulation of NH_4_^+^ **(Fig. 2 C and D; Fig. 4 A-E)**. NADH-GDH plays a unique role from GS-GOGAT in N recycling (Masclaux-Daubresse et al., 2006) and has been widely reported that GDH responds positively to abiotic stress and is important for the detoxification of NH_4_^+^ under stress conditions (Boussama et al., 1999; Bittsánszky et al., 2015; Zhong et al., 2017). The significant increase in GDH activity in *chl1-1* and *chl1-5* in (NH_4_)_2_SO_4_ conditions could diminish the accumulation of NH_4_^+^ **(Fig. 4C)**, and our results reveal that the higher N assimilation capacity in the *chl1-1* and *chl1-5* mutants plays a vital role in mitigating NH_4_^+^ toxicity.

Taken together, our results reveal that enhanced NH_4_^+^ uptake combined with reduced NH_4_^+^ assimilation causes higher NH_4_^+^ accumulation in Col-0 than in the *NRT1.1* mutant lines under high concentrations of (NH_4_)_2_SO_4_, indicating that NH_4_^+^ uptake and assimilation are tightly linked. It has been well documented that Gln is the signal for the regulation of NH_4_^+^ uptake and assimilation (Tabuchi et al., 2007). A previous study revealed that the expression of the cytosolic isoform of GS, *AtGLN1;2*, was strongly induced in roots by NH_4_^+^ and contributed to NH_4_^+^ assimilated in the root (Konishi et al., 2017). Furthermore, the different NH_4_^+^ uptake and assimilation observed in Col-0 and *NRT1.1* mutants suggest that NRT1.1 plays a significant role in regulating these processes. The interaction between *NRT1.1* and *AMT1* genes has been implicated at a physiological level to facilitate N uptake in plants grown in mixed N sources that contain NO_3_^−^ and NH_4_^+^ (Hachiya and Sakakibara, 2017). It is hypothesized that NRT1.1 could affect AMT1s by regulating N assimilation enzymes, as it has been reported that NRT1.1 regulates the expression levels of many N assimilation pathway genes (Bouguyon et al., 2015). The substantial reduction in GS and GOGAT activity in Col-0 plants grown in (NH_4_)_2_SO_4_ **(Fig. 4A and B)** suggests that the pathway for N assimilation enzymes regulated by NRT1.1 is perturbed by high concentration of NH_4_^+^.

Numerous studies have reported the combination of ethylene and NH_4_^+^ accumulation in the development of NH_4_^+^ toxicity symptoms in plants (You and Barker, 2005; Li et al., 2011; Li et al., 2013). Ethylene production could be the link between NH_4_^+^ accumulation and NH_4_^+^ toxic symptoms, as the inhibition of ethylene production ameliorated the symptoms of NH_4_^+^ toxicity even when NH_4_^+^ concentrations in leaves are still high (Li et al., 2013). In our study, when the ethylene mutant *ein2* was grown in (NH_4_)_2_SO_4_ medium, it had a comparable level of NH_4_^+^ concentration to Col-0; furthermore, the growth of Col-0 was unaffected when the plants were grown in (NH_4_)_2_SO_4_ medium and treated with the ethylene inhibitor AVG, whereas ACC induced chlorosis even in *chl1-1* and *chl1-5* plants **(Fig 7A and D)**. These results indicate that NH_4_^+^ accumulation induced ethylene synthesis, which is involved in the development of NH_4_^+^ toxicity in plants. In addition, (NH_4_)_2_SO_4_ up-regulated the expression of genes related to plant senescence in Col-0 plants but not in *chl1-1* and *chl1-5* mutant lines **(Figure S5)**, suggesting the symptoms of NH_4_^+^ toxicity resulted from ethylene induced plant senescence.

## Conclusions

In conclusion, our results suggested that an increase in NH_4_^+^ uptake and reduction in capacity of NH_4_^+^ metabolism is NRT1.1-dependent and the main factor responsible for NH_4_^+^ toxicity **(Fig. 8A and B)**. In wild-type, *NRT1.1* induced higher expression of *AMT1s* and increased NH_4_^+^ uptake in the presence of high concentrations of a sole NH_4_^+^ source. Nevertheless, the ability for NH_4_^+^ assimilation decreased when the plants were grown under high concentrations of sole NH_4_^+^ source medium. Furthermore, with sole NH_4_^+^ source, C and N metabolism was uncoupled, causing an accumulation of soluble sugars and diminished amino acid metabolism. NH_4_^+^ accumulation results in the increased production of ethylene and subsequently higher expression of genes related to plant senescence, and finally NH_4_^+^ toxicity. **(Fig. 8A)**. In the *NRT1.1* mutants *chl1-1* and *chl1-5*, the expression of *AMT1s* was less than that of Col-0. In addition, the C and N metabolism in *nrt1.1* plants were less affected with sole NH_4_^+^ source, and NH_4_^+^ assimilation by the GDH pathway and amino acid synthesis by GOT ensured reduction in the adverse effects of NH_4_^+^. Ultimately, chlorosis did not occur in *NRT1.1* mutants **(Fig. 8B)**. At present, the interaction between nitrate and ammonium transport is a complex and attractive research area. To further explore the regulatory pathways that involve NRT1.1 during NH_4_^+^ uptake, transport, and assimilation are of great importance to elucidate the mechanism of NH_4_^+^ toxicity and to develop methods for managing NH_4_^+^ toxicity.

**Fig. 8.**
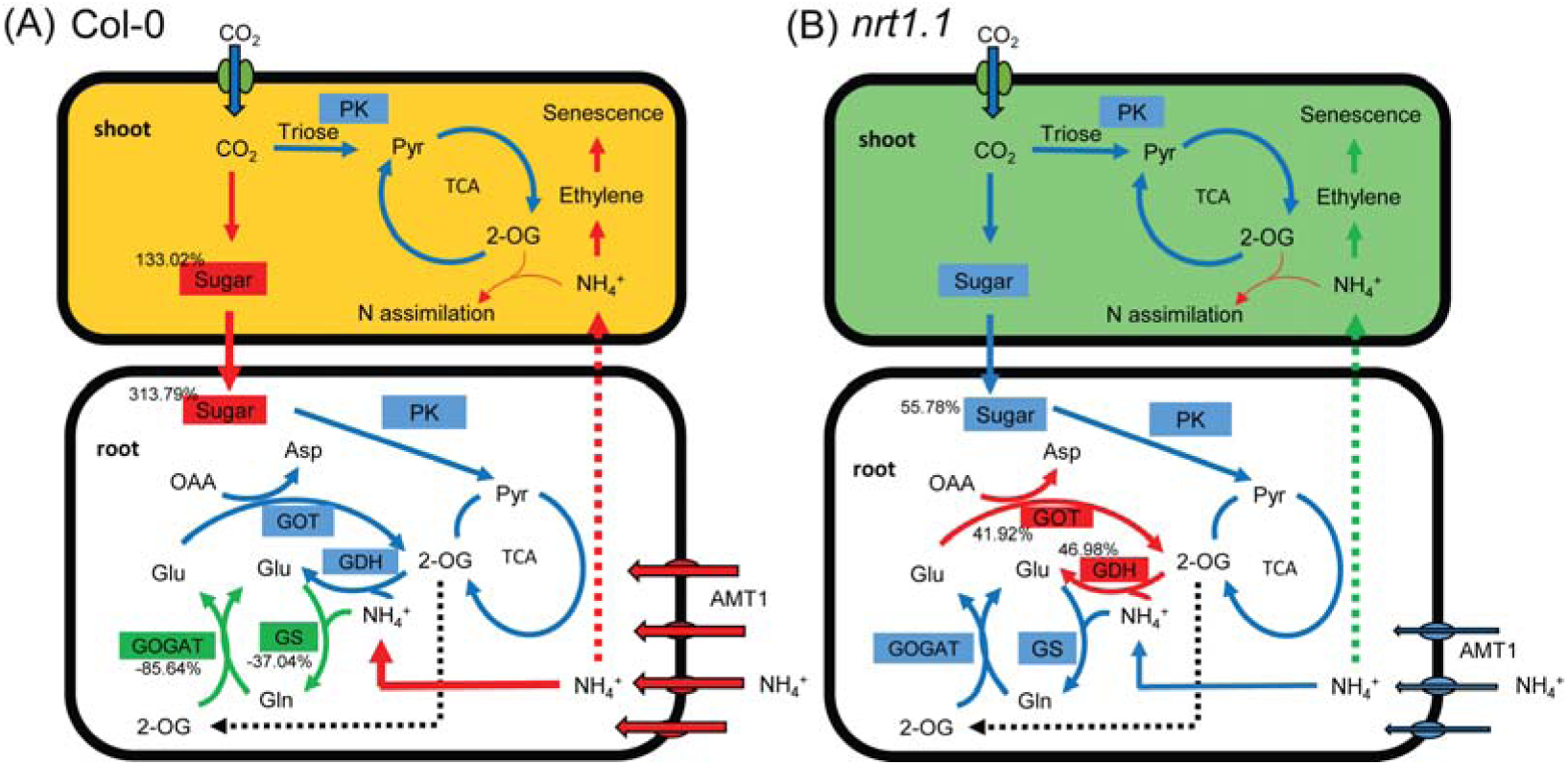
A proposed model of NH_4_^+^ metabolism in Col-0 **(A)** and *NRT1.1* mutants **(B)** grown under high concentrations of a sole NH_4_^+^ source (10 mM in this study). The colors of the rectangles indicate: red, the material or physiological progress was increased; blue, the material or physiological progress was not significantly changed; and green, the material or physiological progress was decreased. Red or green dotted arrows represent NH_4_^+^ flow from roots to shoots speculated to be higher or lower. Red or green solid arrows represent the physiological progress getting stronger or weaker; the numbers nearest to the physiological process indicates the percentage of change in this progress. Compared to plants grown under control conditions, *NRT1.1* induces up-regulated expression of *AMT1 genes* in roots and facilitates NH_4_^+^ uptake. Simultaneously, glutamine synthetase (GS) and glutamate synthetase (GOGAT) activities are inhibited by sole NH_4_^+^ source. Furthermore, sugars accumulate and are stored in roots, leading to the perturbation of carbon and ammonium metabolism. As a result, free NH_4_^+^ is higher and can be transported to the shoot. High NH_4_^+^ concentrations in the shoot induce ethylene synthesis and subsequently cause plant senescence. In *NRT1.1* mutants, the NH_4_^+^ uptake, GS-GOGAT cycle, and carbon and ammonium metabolism are not affected by sole NH_4_^+^ source; in addition, the activity of NADH-dependent glutamate dehydrogenase (GDH) and glutamate-oxaloacetate transaminase (GOT) are enhanced in sole NH_4_^+^ condition, which diminished NH_4_^+^ accumulation and improved amino acid metabolism. Thus, *NRT1.1* mutant plants can better survive environments with high concentrations of a sole NH_4_^+^ source than does the wild type.

## Supplementary information

**Supplementary Table S1** Primers for genes used in this study.

**Supplementary Fig. S1** Effects of (NH_4_)_2_SO_4_ on the phenotype **(A)** and chlorophyll concentration **(B)** of Col-0 and *chl1-9*, and the stomatal conductance **(C)** of Col-0, *chl1-1*, and *chl1-5* plants.

**Supplementary Fig. S2** Comparison of pH values of growth media.

**Supplementary Fig. S3** Effects of (NH_4_)_2_SO_4_ on activity of NH_4_^+^ assimilation enzymes and transaminase in the shoots of Col-0, *chl1-1*, and *chl1-5* plants.

**Supplementary Fig. S4** Effects of (NH_4_)_2_SO_4_ on Thr **(A)**, Met **(B)**, Ile **(C)**, and Ala **(D)** content in roots and shoots of Col-0 and *chl1-1* plants.

**Supplementary Fig. S5** Effects of (NH_4_)_2_SO_4_ on the relative expression of genes related to senescence in Col-0, *chl1-1*, and *chl1-5* plants.

## Acknowledgements

This study was financially supported in part by the National Key R&D Program of China (2017YFD0200103); National Natural Science Foundation of China (Grant No.31101596, 31372130); Hunan Provincial Recruitment Program of Foreign Experts; and the National Oilseed Rape Production Technology System of China; “2011 Plan” supported by The Chinese Ministry of Education; Research and Innovation Project of postgraduates in Hunan province (CX2015B242).

## Abbreviations

*AMT*: ammonium transporter family
*CHL1/NRT1.1/NPF6.3*: nitrate transporter 1/peptide transporter family
GS: glutamine synthetase
GOGAT: glutamate synthetase
GDH: glutamate dehydrogenase
GOT: glutamate-oxaloacetate transaminase
GPT: glutamate-pyruvic transaminase
PPFD: photosynthetic photon flux intensity
PK: pyruvate kinase.

